# Transcriptome Sequencing Reveals Widespread Gene-Gene and Gene-Environment Interactions

**DOI:** 10.1101/010546

**Authors:** Alfonso Buil, Andrew A. Brown, Tuuli Lappalainen, Ana Viñuela, Matthew N. Davies, H. F. Zheng, J. B. Richards, Kerrin S. Small, Richard Durbin, Timothy D. Spector, Emmanouil T. Dermitzakis

## Abstract

Understanding the genetic architecture of gene expression is an intermediate step to understand the genetic architecture of complex diseases. RNA-seq technologies have improved the quantification of gene expression and allow to measure allelic specific expression (ASE)^1-3^. ASE is hypothesized to result from the direct effect of cis regulatory variants, but a proper estimation of the causes of ASE has not been performed to date. In this study we take advantage of a sample of twins to measure the relative contribution of genetic and environmental effects on ASE and we found substantial effects of gene x gene (GxG) and gene x environment (GxE) interactions. We propose a model where ASE requires genetic variability in cis, a difference in the sequence of both alleles, but the magnitude of the ASE effect depends on trans genetic and environmental factors that interact with the cis genetic variants. We uncover large GxG and GxE effects on gene expression and likely complex phenotypes that currently remain elusive.

---

Gene expression is a cellular phenotype that informs about the functional state of the cell. Gene expression by itself is a complex trait that depends on genetic and environmental causes. Many researchers have studied the genetics of gene expression and thousands of expression quantitative loci (eQTL) have been identified within different populations and tissues^4-6^. More recently, the use of RNAseq technologies to measure gene expression has allowed the estimation of allelic specific expression (ASE). ASE measures the difference in expression of two haplotypes of an individual at a specific genetic locus ^1-3^ (Supplementary Fig. 1-A). While eQTLs are population based measures of the effect of genetics on gene expression, ASE is a more direct measure of how gene expression changes at the individual level. In addition, ASE is much less sensitive to technical parameters since such effects would affect both alleles equally. While ASE may occur in a stochastic way within each single cell, measurements from a population of cells for each individual represent the average behavior of the two alleles and, theoretically, are expected to result from the direct effect of genetic regulatory variants in cis. In this study we dissect the underlying causes of ASE, by measuring the relative contribution of genetic and environmental factors and propose biological models of ASE action. To achieve these goals we sequenced the mRNA fraction of the transcriptome of ~400 female twin pairs (~800 individuals) from the TwinUK cohort in four tissues: fat, skin, blood and lymphoblastoid cell lines (LCLs) using 49bp paired-end sequencing in an Illumina HiSeq2000. We sequenced 766 fat samples, 814 LCL samples, 716 skin samples and 384 blood samples and obtained 28M exonic reads per sample on average. Samples were imputed into the 1000 Genomes Phase 1 reference panel. By exploiting the twin structure we can quantify the proportion of variation in ASE that is due to distinct genetic and non-genetic causes.

We used the RNAseq data to estimate ASE in our samples. For every individual at every transcribed heterozygous site we counted the number of reads for which each allele was present. We defined the ASE ratio as the ratio of the logarithm of the counts for the reference allele and the logarithm of the total number of counts. After applying rigorous filtering to control for mapping bias ^1^ (see methods) we called significant ASE sites using a binomial test. At an FDR of 10% we found an average of 303 significant ASE sites in fat, 435 in LCLs, 331 in skin and 141 in blood per individual. To estimate the effect of cis eQTLs on ASE we called cis eQTLs in the four tissues separately. We obtained 9166 cis eQTLs in fat, 9551 in LCLs, 8731 in skin and 5313 in blood (see Methods).

To quantify genetic and environmental sources of variation in ASE we developed an extension of the classical variance components approach based on the correlations within MZ and DZ twin pairs. For genes with at least one eQTL, we looked at population level variation in the ASE ratio (treated as a quantitative phenotype), for those sites with an appropriate ASE measurement. We estimated the correlation of this phenotype within MZ and DZ twins and observed that the correlation among DZs was higher than half of that of the MZs (Figure 1). That could indicate a potential shared environment component but, in our case, it is more likely due to the fact that the cis eQTL has a large effect on ASE and our DZ twins are genetically more similar than random mating predicts at the ASE locus (mean Identical By Descent (IBS) coefficient at the eQTL for DZ twins is 0.9). Indeed, when we looked at correlation between DZ twins that are IBS=.5 (and hence share half of the contribution of the additive eQTL of MZ twins) we observed that this correlation is less than half of the correlation between MZ twins (Figure 1), indicating the potential presence of non additive genetic effects.

**Figure 1.**
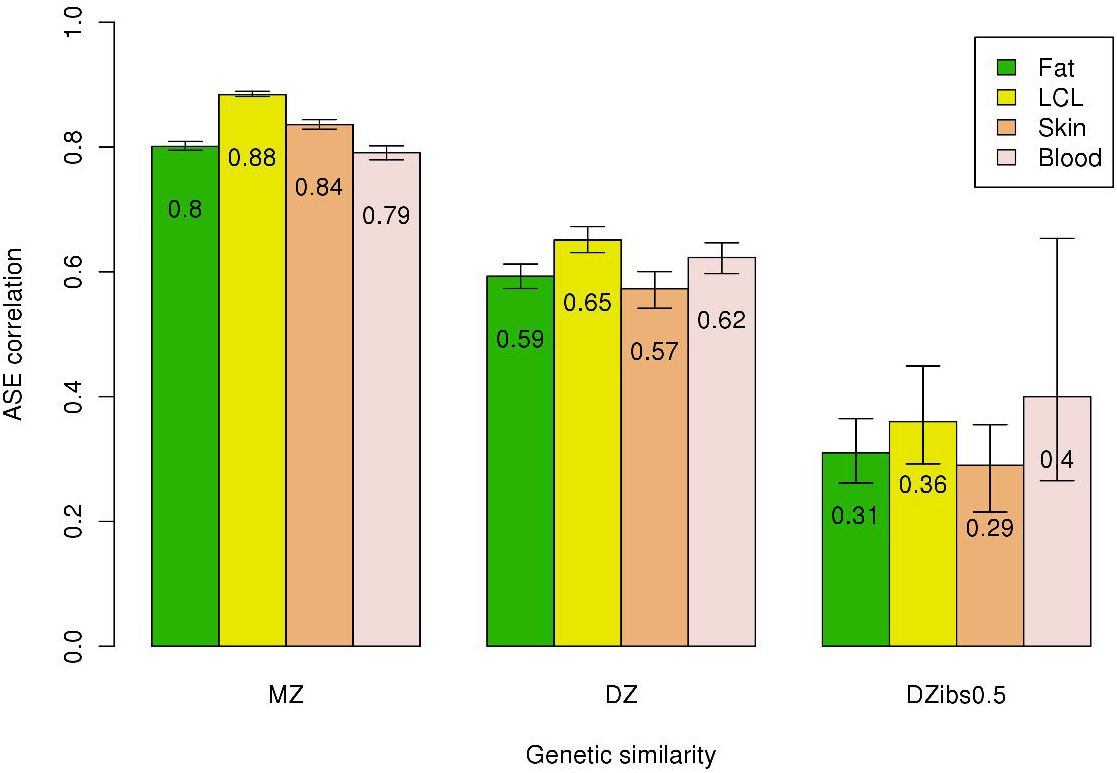
ASE correlation among twin pairs for different categories of genetic similarity. MZ: monozygotic twins; DZ: dizygotic twins; DZibs05: DZ twins with identical by state (IBS) equal to 0.5 at the eQTL locus. The 95% confidence intervals were calculated by using 1000 bootstrap permutations.

To incorporate these complexities in the model, we separated the twin pairs depending on the average genetic similarity genome wide (1 for MZ, 0.5 for DZ), genetic similarity at the locus based on the Identity By Descent (IBD) status in the cis region surrounding the gene, and the genetic similarity of the eQTL based on the IBS status at the eQTL locus. We estimated the correlations of the ASE ratio for each category of twins and modeled these correlations as a function of six variance components (see methods). These components represent the proportions of variance in ASE that could be explained by environmental variation, by the principal eQTL, by other variants in cis, by variants in trans, and by genetic interactions. As recombination is unlikely to have occurred within the cis window, cis epistatic interactions are generally not broken up within twin pairs and thus their contribution to variance is effectively additive. Instead we looked at calculating the proportion of variance explained by cis-trans interactions. We found that the heritability (the sum of all the genetic components, eQTL + cis + trans + interactions) of ASE ranges from 62% to 88% (Figure 2). The effect of the best cis eQTL per gene accounts for 26% to 46% of the variance on ASE. That means that nearly half of the heritability of ASE is due to a common cis eQTL. The remaining is due to other genetic effects in cis (11% to 22% of the ASE variance) and genetic interaction effects (11% to 29% of the ASE variance). We did not observe significant additive trans effects. We found a significant effect of the shared environment only in blood (11% of the ASE variance). That could be due to the fact that blood is more heterogeneous than the other tissues, with variable proportions of different cell types in individuals and shared environment affecting the counts of different cell types. In the shared environment component we are likely picking up cell-type specific effects. We used 1000 bootstrap permutations to calculate confidence intervals of our variance components estimates (Figure 2). In summary, the main causes of ASE are genetic variants in cis, as expected, but between 38% and 49% of the variance in the ASE ratio is due to genetic interactions and environmental factors.

**Figure 2.**
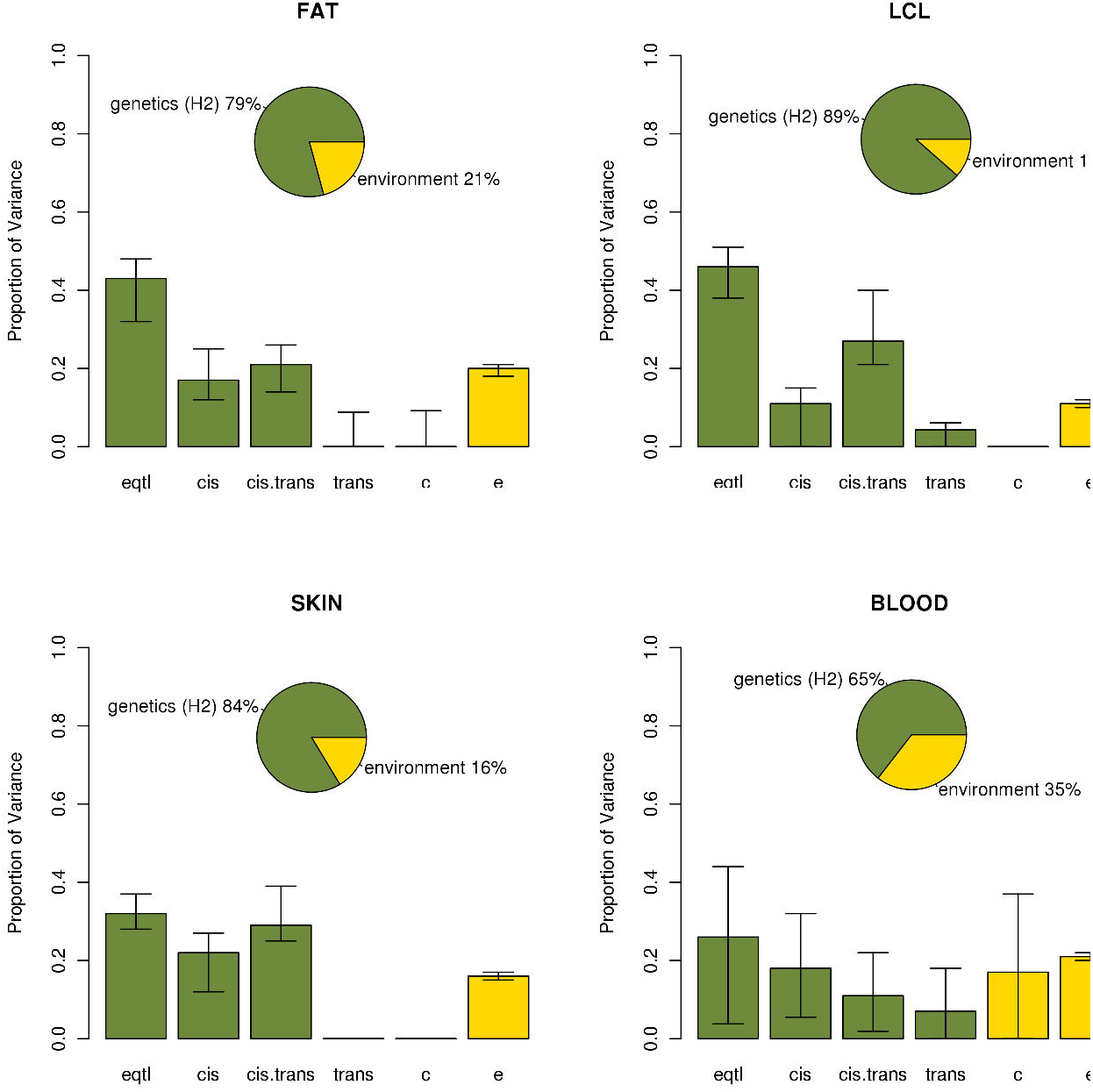
Variance components for ASE for the model with cis x trans interaction: ‘H2’ (equal to eqtl + cis + cis.trans + trans) is the heritability of ASE, ‘eqtl’ is the proportion of variance explained by a common eQTL, ‘cis’ is the proportion of variance explained by other variants in cis, ‘cis.trans’ is the proportion of variance explained by interactions between cis and trans genetic variants, ‘trans’ in the proportion of variance explained by genetic variants in trans, ‘c’ is the proportion of variance explained by the shared environment and ‘e’ is the proportion of variance explained by the individual environment. The 95% confidence intervals where calculated using 1000 bootstrap permutations.

To confirm that the interaction component is due to cis x trans interactions we adjusted a model without the interaction component and we observed that the interaction effects moved mainly to the trans component (Supplementary Fig. 2). This result strongly suggests that the effect of the genetic regulatory variants in cis that cause ASE is modulated to a substantial extent by genetic variants in trans.

Our variance components model shows that genetic effects do not explain all the observed variance in ASE and that environmental factors can have an effect on ASE. Given the nature of the ASE, these environmental effects should be mainly mediated by true biological mechanisms mediated by epigenetic mechanisms and much less likely by technical and experimental effects. Environmental/epigenetic effects alone cannot create allelic imbalance as this is inferred by a population of cells; to observe ASE a cis DNA sequence effect is required. We therefore postulated the existence of gene x environment (GxE) interactions affecting ASE. To identify cases of GxE interactions we used the distance (degree of discordance) of ASE in MZ twins. Previous studies have used MZ twin pairs discordant for a phenotype to map genetic variants affecting that phenotype ^7-9^. In this case, our phenotype is the ASE ratio and, for every site, we calculated the association between distance ASE between individuals of an MZ pair and SNPs around the site. We found evidence of GxE in fat and LCLs but almost nothing for skin and blood (Supplementary Tables 1-4). One of the top hits in LCLs is the Epstein-Barr virus induced 3 gene (EBI3) (Figure 3). That means that ASE at the *EBI3* gene depends on the interaction of cis genetic variants and an environmental factor that, likely, in this case is the transformation process of the B cells with EBV. The two top hits in fat are *ADIPOQ* and *ACSL1*, two genes that code for Adiponectin and Long-chain-fatty-acid—CoA ligase 1 proteins respectively (Figure 3). These two proteins are functionally related: both participate in the gene ontology biological processes ‘response to fatty acid’ and ‘response to nutrient’, and both are known to be regulated by environmental factors such as diet and exercise in a genotype dependent manner ^10-13^. Attempts to link it to environmentally affected phenotypes (BMI, glucose levels, insulin levels) did not show any signal, which is not surprising since these are phenotypes affected by the environment and not direct environmental measures. The analysis above suggests the presence of GxE interactions on gene expression.

**Figure 3.**
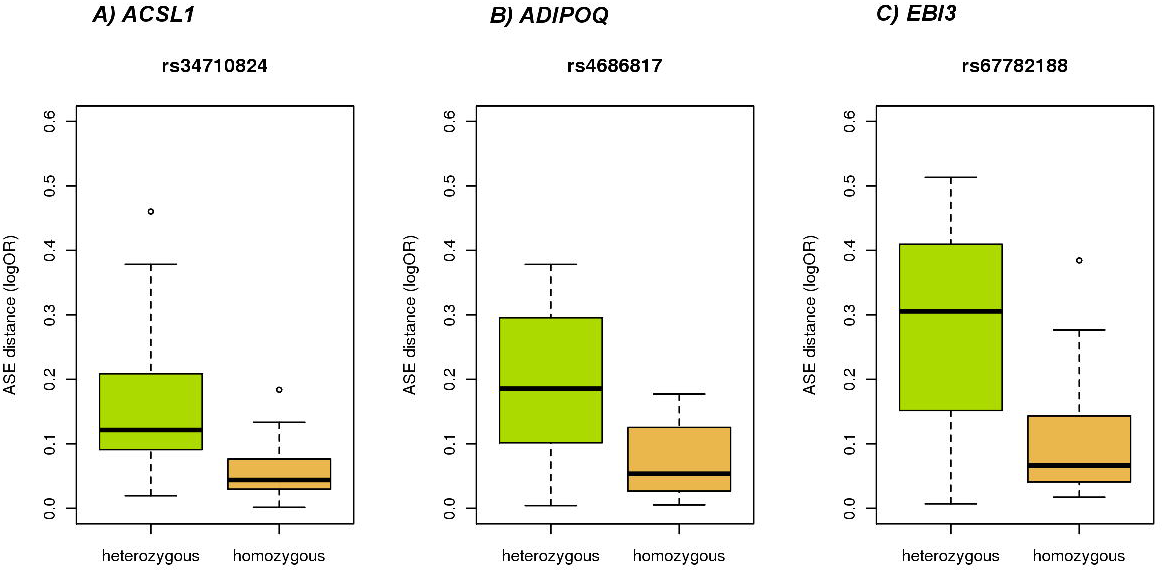
GxE examples discovered using discordant MZ twins analysis. MZ twin pairs show a different ASE effect on some genes depending on the genotype of specific SNPs. The Y axis shows the ASE difference between MZ twins. Since MZ twins are genetically identical, this association reflects the interaction of the SNP with an unknown environment. A) ASE in gene ACSL1 shows GxE interaction with SNP r334710824 in Fat; B) ASE in gene ADIPOQ shows GxE interaction with SNP rs4686817 in Fat; C) ASE in gene EBI3 shows GxE interaction with SNP rs67782188 in LCLs.

In conclusion, these results show a complex genetic architecture for cis-regulation of gene expression measured through ASE. We propose a model where the allelic imbalance in expression (ASE) requires genetic variability in cis, however, the magnitude of the ASE effect depends on trans genetic and environmental factors that interact with the cis genetic variants (Supplementary 1B, 1C). About 38% to 49% of the variance of the observed ASE is not explained by additive genetic effects. This means that a substantial amount of the variance observed on ASE, and therefore in regulation of gene expression, is due to GxG and GxE interactions. It is worth to note that our results show no additive trans effects on ASE. That does not mean that there are not additive trans effects affecting gene expression; that means that the trans effects affecting ASE are not additive. We found an example of GxE interaction on gene expression that has been widely described in the literature (Adiponectin) supporting the validity of our approach. However, the limitation on power due to the sample size prevented us to discover specific associations in most other cases. Allelic gene expression is the molecular phenotype closest to the action of genetic variation. The presence of widespread GxG and GxE interactions affecting this phenotype implies that GxG and GxE can be important in other complex phenotypes, including diseases. The proposed model has implications for the interpretation of the effect of GWAS genetic variants on complex diseases. About 80% of the GWAS signals are estimated to be regulatory variants. The search for GxG and GxE interactions conditioning on relevant biological models rather than whole genome anonymous searches is likely to recover a substantial fraction of genetic and non-genetic variance associated with disease risk.

## Acknowledgements

This work has been funded by the EU FP7 grant EuroBATS (No. 259749) which also supports AAB, AB, MND, AV, TDS. AAB is also supported by a grant from the South-Eastern Norway Health Authority, (No. 2011060). RD is supported by the Wellcome Trust (No. 098051). The Louis-Jeantet Foundation, Swiss National Science Foundation, Eurpean Research Council and the NIH-NIMH GTEx grant supports ETD. TS is an NIHR senior Investigator and holder of an ERC Advanced Principal Investigator award. JBR and HZ are supported by the Canadian Institutes of Health Research, Fonds de Recherche Sante du Quebec, and the Quebec Consortium for Drug Discovery. Most computations were performed at the Vital-IT (http://www.vital-it.ch) Center for high-performance computing of the SIB Swiss Institute of Bioinformatics. The TwinsUK study was funded by the Wellcome Trust; EC FP7(2007-2013) and National Institute of Health Research (NIHR). SNP Genotyping was performed by The Wellcome Trust Sanger Institute and National Eye Institute via NIH/CIDR. We thank the twins for their voluntary contribution to this project. RNAseq and Genotyping data are available under EGA accesion numbers XXXXXXX and YYYYYYY (to be provided at a later stage).

## Online Methods

### Genotying and imputation

Samples were genotyped on a combination of the HumanHap300, HumanHap610Q, 1M-Duo and 1.2MDuo 1M Illumnia arrays. Samples were imputed into the 1000 Genomes Phase 1 reference panel (data freeze, 10/11/2010)^14^ using IMPUTE2^15^ and filtered (MAF<0.01, IMPUTE info value < 0.8).

### RNA processing

Samples were prepared for sequencing with the IlluminaTruSeq sample preparation kit (Illumina, San Diego, CA) according tomanufacturer’s instructions and were sequenced on a HiSeq2000 machine. Afterwards, the 49-bp sequenced paired-end reads were mapped to the GRCh37reference genome^16^ with BWA v0.5.9^17^. We use genes defined as protein coding in the GENCODE 10 annotation^18^.

### Quantification of allelic specific expression

We defined the ASE ratio as the ratio of the logarithm of the counts for the reference allele and the logarithm of the total number of counts. We called significant ASE sites using a binomial test. We exclude sites that are susceptible to allelic mapping bias: 1) sites with 50bp mapability < 1 based on the UCSC mapability track, implying that the 50bp flanking region of the site is non-unique in the genome, and 2) simulated RNA-seq reads overlapping the site that show >5% difference in the mapping of reads that carry the reference or non-reference allele ^1^. Finally, to correct for systematic bias in allelic ratios we calculated the overall reference to total allele ratio for each individual for each SNP base combination. These ratios were then used as the expected ratios in the binomial test.

### eQTL discovery

Since our data samples are twins, they are not independent observations and we have to take that into account in our models. To find cis eQTL we used the two steps strategy proposed in Svishcheva et al. ^19^. First, we kept the residuals of a mixed model to remove the effects of the family structure using the implementation in GenAbel R packate ^20^. In the second step we performed a linear regression of those residuals on the SNPs in a 1Mb window around the transcription start site for each gene, using MatrixeQTL R package ^21^. We assessed statistical significance through 2000 permutations. Previous to the analysis we removed the effects of technical covariates using the factor analysis strategy implemented in PEER ^22^ and transformed the data using a rank normal transformation.

### Variance Components Models

Classical variance components models in twins model the phenotypic correlation between MZ twins and DZ twins as a function of the additive genetic variance and the shared environment variance^23^:

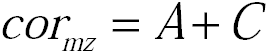

and

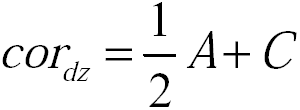

where A represents the additive genetic effects and C the effects due to the common environment between the twin pair (events that affect each twin in a different way). The individual environmental effect (events that occur to one twin but the other) would be *E* = 1 − *cor*_*mz*_. From the two equations above, we get that heritability can be estimated as:

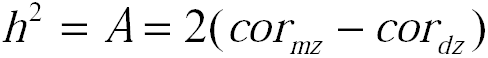

Here, we extend this model to incorporate new sources of variation:

- *A*_*qtl*_: additive effect due to the best eQTL
- *A*_*cis*_: other genetic additive effects in cis
- *A*_*trans*_: additive genetic effects in trans
- *I*: epistatic interaction between trans and cis genetic effects

Where *A* = *A*_*qtl*_ + *A*_*cis*_ + *A*_*trans*_ + *I*

Then, our model has six variance component: 1) variance due to the effect of the major ciseQTL (the IBS status at this locus), 2) variance due to the rest of the genetic variants in cis (including the effect of rare variants, captured by the IBD status), 3) variance due to genetic variants in trans (the genome-wide IBD), 4) variance due to non-additive genetic effects (genetic interactions), 5) variance due to the shared environmental effect and 6) variance due to the individual environmental effect.

The equations of the extended model are:

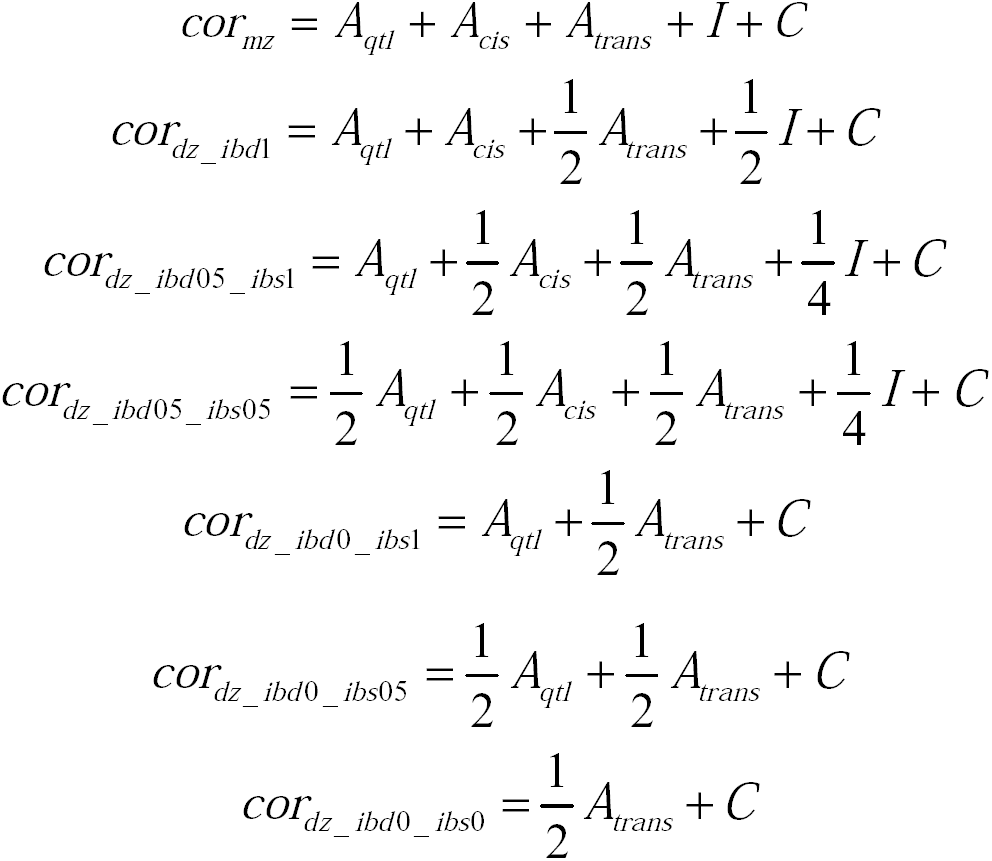

Where:

*cor*_*mz*_ is the correlation within MZ twins
*cor*_*dz_ibd*1_ is the correlation within DZ twins that are IBD=1 at the gene
*cor*_*dz_ibd* 05_*ibs*1_ is the correlation within DZ twins that are IBD=0.5 at the gene and IBS=1 at the eQTL
*cor*_*dz_ibd* 05_*ibs* 05_ is the correlation within DZ twins that are IBD=0.5 at the gene and IBS=0.5 at the eQTL
*cor*_*dz_ibd* 0_*ibs*1_ is the correlation within DZ twins that are IBD=0 at the gene and IBS=1 at the eQTL
*cor*_*dz_ibd* 0_*ibs* 05_ is the correlation within DZ twins that are IBD=0 at the gene and IBS=0.5 at the eQTL
*cor*_*dz_ibd* 0_*ibs* 0_ is the correlation within DZ twins that are IBD=0 at the gene and IBS=0 at the eQTL

To calculate these correlations we used sites covered by at least 30 reads, showing significant ASE in genes with at least one cis eQTL. Since the number of individuals that have ASE at a given site is small, we analyzed all the sites together to get a global estimate of the variance components. This strategy has been used previously with gene expression data^4,24^.

To solve the system of equations we used the non-linear optimization package Rsolnp from the R statistical environment^25^. We estimated the solution that minimizes the quadratic errors, forcing the variances components to be positive.

### Genotype by Environment Interaction

We used a discordant twin analysis where the difference in the ASE ratio between MZ twins at a given gene is regressed on the SNPs in a 1Kb window around the transcription start site of the gene. Since we are looking for an effect on ASE we expect a similar behavior for the two homozygous genotypes. Then, for the association analysis we coded the genotypes in two categories: homozygous and heterozygous.

